# First isolation and characterisation of Alongshan virus in Russia

**DOI:** 10.1101/862573

**Authors:** Ivan S. Kholodilov, Alexander G. Litov, Alexander S. Klimentov, Oxana A. Belova, Alexandra E. Polienko, Nikolai A. Nikitin, Alexey M. Shchetinin, Anna Y. Ivannikova, Lesley Bell-Sakyi, Alexander S. Yakovlev, Sergey V. Bugmyrin, Liubov A. Bespyatova, Larissa V. Gmyl, Svetlana V. Luchinina, Anatoly P. Gmyl, Vladimir A. Gushchin, Galina G. Karganova

**Author notes:** These authors contributed equally to this article (Ivan S. Kholodilov, Alexander G. Litov). Deceased (Anatoly P. Gmyl). Corresponding author: Dr. Galina G. Karganova; “Chumakov Institute of Poliomyelitis and Viral Encephalitides” FSBSI “Chumakov FSC R&D IBP RAS”, prem. 8, k.17, pos. Institut Poliomyelita, poselenie Moskovskiy, Moscow, 108819, Russia.

## Abstract

In recent decades, many new flavi-like viruses have been discovered predominantly in different invertebrates and, as was recently shown, some of them may cause disease in humans. The Jingmen tick virus (JMTV) group holds a special place among flavi-like viruses because, in contrast to the “classic” flaviviruses and other flavi-like viruses, they have a segmented ssRNA(+) genome. Two segments of the JMTV genome have homology with regions of the flavivirus genome encoding polymerase and helicase-protease proteins. JMTVs have several open reading frames (ORF) in segments encoding glycoprotein(s) and capsid protein and these ORF are specific only to them. JMTVs greatly differ in virion size.

We isolated three strains of Alongshan virus (ALSV), which is a representative of the JMTV group, from adult *Ixodes persulcatus* ticks collected in two geographically-separated Russian regions in the tick cell line IRE/CTVM19. One of the strains persisted in the IRE/CTVM19 cells without cytopathic effect for three years. Most virions purified from tick cells were spherical with a diameter of approximately 40.5 nm. In addition, we found smaller particles of approximately 13.1 nm in diameter. We obtained full genome sequences of all four segments of two of the isolated ALSV strains, and partial sequences of one segment from the third strain. Phylogenetic analysis on genome segment 2 of the JMTV group clustered our novel strains with other ALSV strains. We found evidence for the existence of a novel upstream ORF in the glycoprotein-coding segment of ALSV and other members of the JMTV group.

**Significance Statement:** We isolated three strains of Alongshan virus (ALSV) from adult *Ixodes persulcatus* ticks from two geographically separate areas of Russia in the *Ixodes ricinus* tick cell line IRE/CTVM19. One of the strains persisted in the IRE/CTVM19 cells without cytopathic effect for three years. Our study confirmed the value of tick cell lines in virus isolation and maintenance of persistent infection. The majority of virions of the ALSV strain Miass527 were enveloped spherical particles with a diameter of 40.5±3.7 nm. We found evidence for the existence of a novel upstream ORF in the glycoprotein-coding segment of ALSV and other members of the Jingmen tick virus group.

## Introduction

In recent decades, many new flavi-like viruses have been discovered in different invertebrates. All of them share homology in the nonstructural proteins with the well-studied flavivirus RNA-dependent RNA polymerase NS5 and RNA helicase-protease NS3, but they differ greatly from each other regarding other proteins, as well as in overall genome size and structure (Shi et al., 2016).

The Jingmen tick virus (JMTV) group holds a special place among flavi-like viruses because in contrast to the “classic” flaviviruses and other flavi-like viruses, they have a segmented ssRNA(+) genome (Ladner et al., 2016; Maruyama et al., 2014; Qin et al., 2014; Shi et al., 2016; Wang et al., 2019a), which is more commonly observed in viruses of fungi and plants (Lefkowitz et al., 2018). The only other known viruses with a segmented ssRNA(+) genome are representatives of the family Nodaviridae and infect insects (*Alphanodavirus*) and fish (*Betanodavirus*) (Yong et al., 2017).

Two segments of the JMTV group genome are homologous to the regions of the Flavivirus genome encoding NS3 and NS5 proteins. The remaining segments are specific only to JMTVs. Additionally, in contrast to the “classic” flaviviruses that have one long open reading frame (ORF) and virion size of 40-60 nm, these viruses have several ORFs within single segments and virion sizes varying from 30-35 nm to 70-100 nm (Ladner et al., 2016; Qin et al., 2014; Wang et al., 2019a). It has also been suggested that genomic RNAs of a multicomponent virus can be packaged in separate virions (Ladner et al., 2016), although this has not yet been demonstrated.

The geographic distribution of the JMTV group is very wide, encompassing Asia (Dinçer et al., 2019; Jia et al., 2019; Qin et al., 2014; Shi et al., 2016), Europe (Dinçer et al., 2019; Emmerich et al., 2018; Kuivanen et al., 2019; Temmam et al., 2019), Central and South America (Ladner et al., 2016; Maruyama et al., 2014; Pascoal et al., 2019; Temmam et al., 2019; Villa et al., 2017) and Africa (Ladner et al., 2016; Temmam et al., 2019). Virus RNAs of this group were detected not only in different species of insects, e.g. mosquitoes (Wang et al., 2019a) and *Drosophila melanogaster* (Webster et al., 2015), but also in the nematode *Toxocara canis* (Callister et al., 2008; Tetteh et al., 1999) and ticks (Dinçer et al., 2019; Qin et al., 2014; Shi et al., 2016) including *Rhipicephalus microplus* (Pascoal et al., 2019; Temmam et al., 2019), *Ixodes ricinus* (Kuivanen et al., 2019; Temmam et al., 2019) and *Ixodes persulcatus* (Jia et al., 2019; Wang et al., 2019a). JMTV were isolated from *Rhipicephalus microplus* ticks with restricted *in vitro* multiplication in tick and mammalian cells (Maruyama et al., 2014; Qin et al., 2014), and from *Amblyomma javanense* ticks with continuous cultivation in a tick cell line (Jia et al., 2019). Some representatives of the JMTV group were detected in patients with other infections transmitted by arthropods (Emmerich et al., 2018).

Alongshan virus (ALSV), a representative of the JMTV group, was detected in *I. persulcatus* ticks, *Culex tritaeniorhynchus* and *Anopheles yatsushiroensis* mosquitoes in China (Wang et al., 2019a) and later in *I. ricinus* ticks in Finland and France (Kuivanen et al., 2019; Temmam et al., 2019). ALSV was suspected to infect mammals and cause illness in humans (Wang et al., 2019b, 2019a).

Our work presents data on the isolation of ALSV in the Russian Federation and new data on the genome organization of this and other known representatives of the JMTV group.

## Results

### Detection and Isolation of Alongshan virus strains

We collected unfed adult *I. persulcatus* ticks in Chelyabinsk region and in the Republic of Karelia in May–June 2014 (Supplementary file 1). The ticks were pooled, homogenised and the homogenates screened for flavivirus NS5 RNA. The PCR products from three flavivirus-positive tick pools were Sanger-sequenced. The resultant sequences were analysed with BLAST. All three isolates, named Miass519, Miass527 and Galozero-14-T20426, were found to be 90.48 – 91.03% similar to Alongshan virus.

To isolate the virus from tick homogenate, we inoculated cultures of the IRE/CTVM19 tick cell line. An IRE/CTVM19 culture inoculated with the Miass527 sample was kept for a year with regular medium changes but without passaging or subculture; individual tick cell cultures can be maintained for long periods without subculture (Bell-Sakyi et al., 2007). Since the virus did not cause any cytopathic effect in these cells, we changed the medium once every 14 days. Once every 1-2 months we checked the culture medium by RT-PCR for presence of the virus, using specific primers MiassF and MiassR (Supplementary file 2). After a year of maintenance, 200μl of supernate from the Miass527-infected IRE/CTVM19 culture was passaged into a new flat-sided culture tube with uninfected IRE/CTVM19 cells. This new Miass527-infected culture was subcultured by splitting the cells at a ratio of 1:1 twice in the first 6 months of infection and after that was kept without passaging or subculture. In this condition Miass527-infected cells have been monitored for over 3 years (observation is still continuing). IRE/CTVM19 cells inoculated with samples Miass519 and Galozero-14-T20426 were each subcultured 1:1 twice in the first 2 months after infection and, at the time of writing, have been monitored for 4 months.

RT-PCR analysis revealed that the strain Miass527 successfully persisted in the tick cell line for a year, followed by one passage and further three years persistence at the time of writing. The other two strains, Miass519 and Galozero-14-T20426, had persisted for at least 4 months at the time of writing.

For two strains, Miass519 and Miass527, high-throughput sequencing of virus purified from IRE/CTVM19 cell supernate was carried out to obtain full genomes, while for the Galozero-14-T20426 strain partial sequences of segment 2 were obtained via Sanger sequencing using primers JMun1S and Miass_gly_1R (Table 1, Supplementary file 2). For the phylogenetic analysis, we used RNA sequences from 18 GenBank entries of representatives of the JMTV group and the three ALSV strains isolated in the present study (Table 1). The phylogenetic analysis was performed on a 1732 bp fragment of segment 2. Interestingly, one of the JMTV isolates (MN095520 – isolate JMTV/I.ricinus/France) clustered with the ALSV sequences. For consistency, we included this JMTV sequence in all our analyses of ALSV (see below). All three of our strains belonged to ALSV and formed one monophyletic group with the strain H3 isolated in China (Figure 1).

**Table 1.**
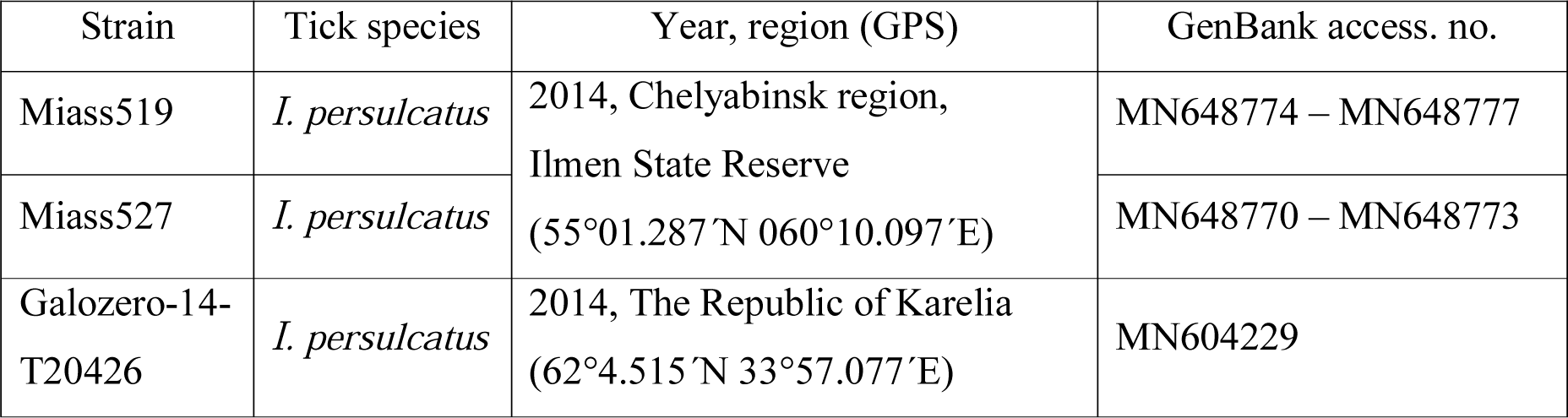
Alongshan virus strains isolated in unfed adult *Ixodes persulcatus* ticks from two geographically separate regions of Russia

| Strain | Tick species | Year, region (GPS) | GenBank access. no. |
| --- | --- | --- | --- |
| Miass519 | *I. persulcatus* | 2014, Chelyabinsk region,<br>Ilmen State Reserve<br>(55°01.287'N 060°10.097'E) | MN648774 – MN648777 |
| Miass527 | *I. persulcatus* |  | MN648770 – MN648773 |
| Galozero-14-T20426 | *I. persulcatus* | 2014, The Republic of Karelia<br>(62°4.515'N 33°57.077'E) | MN604229 |

**Figure 1.**
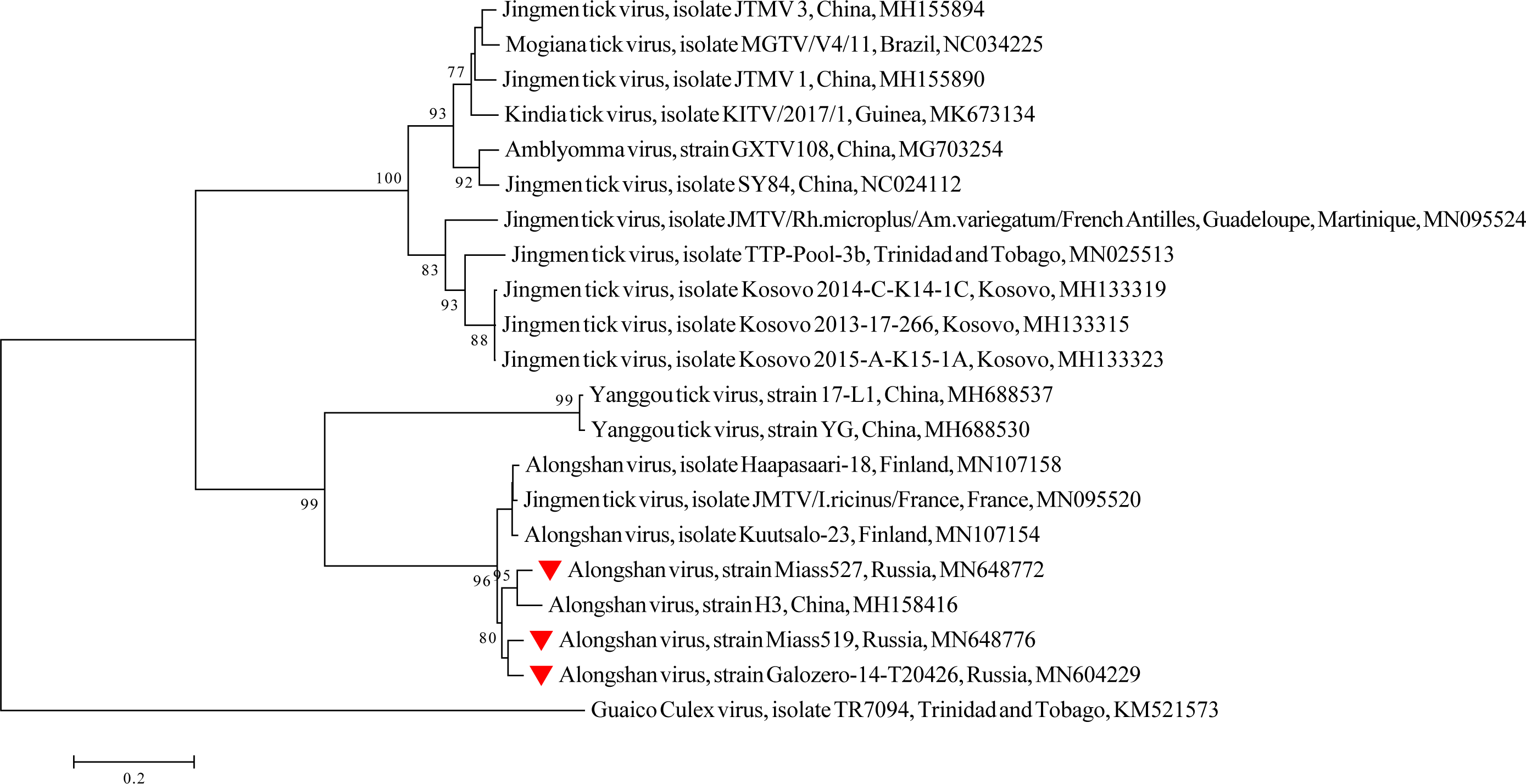
Phylogenetic tree of a 1732 bp fragment of segment 2 of Alongshan virus and other members of the Jingmen tick virus group. Phylogenetic trees were constructed using MEGA 6.0 with the neighbour-joining method (1000 bootstrap replications). Bootstrap values (>70%) are shown at the branches. GenBank accession numbers are listed for each strain. 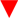 Strains from the present study.

### Transmission Electron Microscopy

We used pool of cell culture supernate collected after one passage of strain Miass527 in IRE/CTVM19 cells, at 17, 19, 20 and 32 months after initial isolation for transmission electron microscopic examination. The IRE/CTVM19 tick cell line was recently reported to contain RNA sequences with similarity to rhabdoviruses (Bell-Sakyi and Attoui, 2016). To prevent confusion between ALSV and this putative rhabdovirus in our microscopic examination, we first screened the pellet obtained after ultracentrifugation of uninfected IRE/CTVM19 for rhabdovirus RNA using specific primers (Supplementary file 2). We detected an unknown rhabdovirus and named it IRE/CTVM19-associated rhabdovirus. Since this new rhabdovirus was also detected in the Miass527-infected IRE/CTVM19 cell pellet, we subjected the dissolved pellet to sucrose density gradient separation and selected the fraction with the largest amount of ALSV RNA and very low levels of rhabdovirus RNA (Supplementary file 3). The microscopy revealed that most virions of the strain Miass527 were enveloped spherical particles with a diameter of 40.5±3.7 nm (Figure 2, A, B, Supplementary file 4). They had either electron-translucent or electron-dense cores. The viral particles with an electron-dense core could have lost genomic RNA (Figure 2, B, Supplementary file 6). In the same gradient fraction we found small spherical particles with a diameter of 13.1±2.1 nm, which had electron-translucent or electron-dense cores (Figure 2, C, D, Supplementary file 5). These small particles could be virions with incomplete genome, protein structures or some other unidentified component of IRE/CTVM19 cells. The pellet obtained after ultracentrifugation of uninfected IRE/CTVM19 culture supernate was separated by sucrose density gradient ultracentrifugation. For transmission electron microscopy we used the same fraction of the gradient as used for the ALSV-infected IRE/CTVM19 cells. In this fraction we found small spherical particles with a diameter of 15.7 ± 1.8 nm, which had electron-dense cores (Figure 2, E, Supplementary file 6). These small particles could be virions of an unknown virus, protein structures or some other unidentified component of IRE/CTVM19 cells.

**Figure 2.**
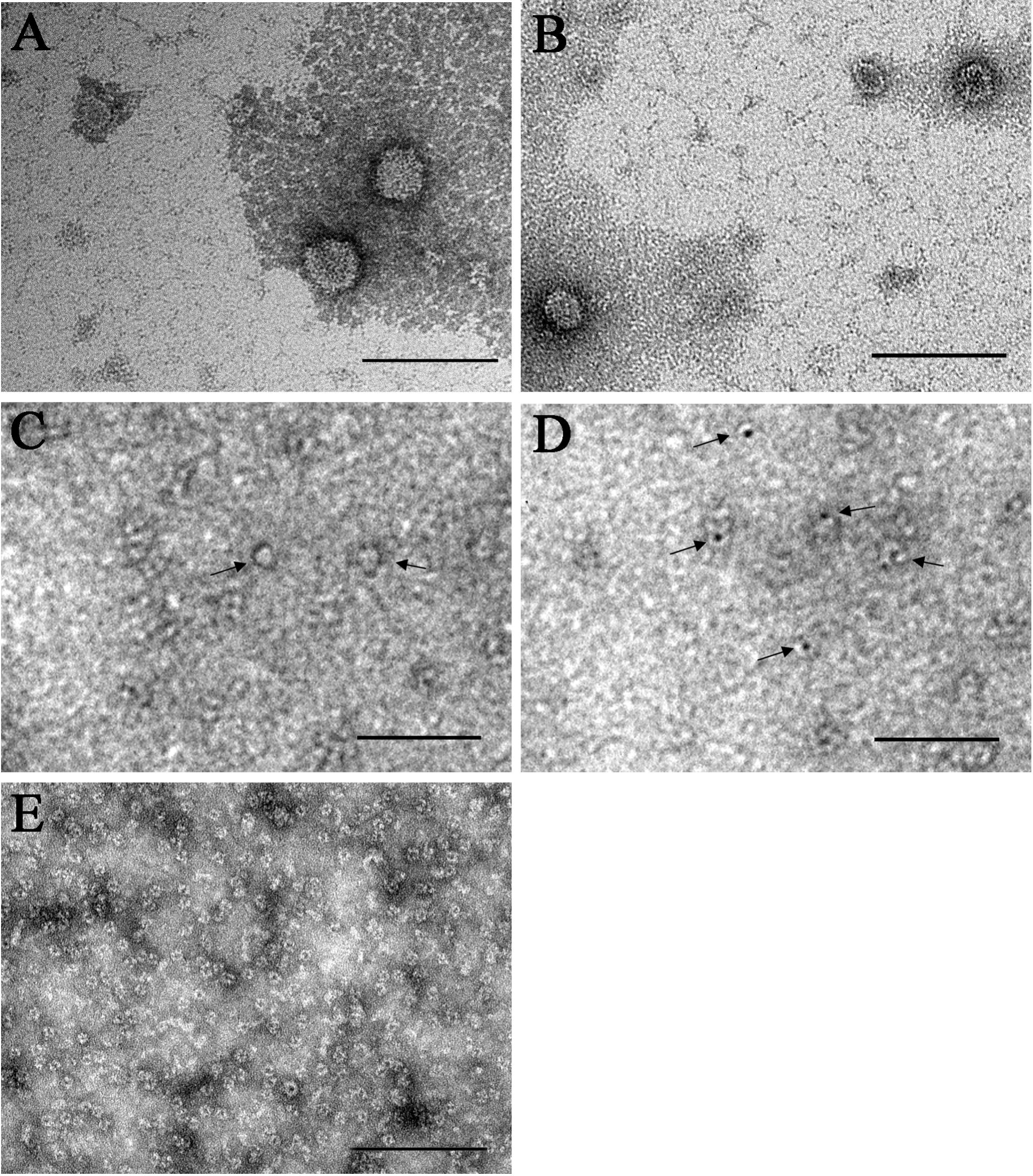
Transmission electron microscopy. Electron micrographs of purified viral particles of the Alongshan virus strain Miass527 propagated in IRE/CTVM19 cells (A, B). Small particles with electron-translucent (C) or electron-dense (D) cores were seen. Small particles with electron-dense cores were also seen in the equivalent fraction of ultracentrifuged supernate from uninfected IRE/CTVM19 cells (E). Samples were stained with 2% uranyl acetate. Scale bars, 100 nm.

### Full genome, genome structure and proposed novel elements

We sequenced full genomes of the strains Miass527 and Miass519. Segments 1, 3 and 4 of the newly-isolated Miass strains were more similar to each other than to previously-described ALSV strains. However, segment 2 of the strain Miass527 was more similar to the Chinese strain H3 than to Miass519, while segment 2 of the strain Miass519 was closer to JMTV/I.ricinus/France than to Miass527 (Table 2).

**Table 2.** Percentage nucleotide identity of the four genome segments of different strains of Alongshan virus for which full genomes were available at the time of writing. Genbank accession numbers for each strain and isolate are as follows: Miass519 (MN648774 – MN648777) (this study); Miass527 (MN648770 – MN648773) (this study); H3 (MH158415 – MH158418) (Wang et al., 2019a); Kuutsalo-23 (MN107153 – MN107155) and Haapasaari-18 (MN107157 – MN107160) (Kuivanen et al., 2019); JMTV/I.ricinus/France (MN095519 – MN095522) (Temmam et al., 2019)

|  | Strain<br>Miass527 | Strain<br>H3 | Strain<br>Kuutsalo-23 | Strain<br>Haapasaari-18 | Isolate<br>JMTV/I.ricinus<br>/France |
| --- | --- | --- | --- | --- | --- |
| **Segment 1. NS5-like protein** |  |  |  |  |  |
| Strain Miass519 | 90.6% | 89.0% | 89.2% | 88.9% | 89.1% |
| Strain Miass527 |  | 89.6% | 89.2% | 88.6% | 89.7% |
| Strain H3 |  |  | 89.6% | 89.2% | 90.1% |
| Strain Kuutsalo-23 |  |  |  | 95.6% | 96.0% |
| Strain Haapasaari-18 |  |  |  |  | 95.9% |
| **Segment 2. VP1a and VP1b** |  |  |  |  |  |
| Strain Miass519 | 92.6% | 92.0% | 93.7% | 93.6% | 94.0% |
| Strain Miass527 |  | 94.7% | 92.4% | 92.2% | 93.0% |
| Strain H3 |  |  | 91.7% | 91.6% | 92.1% |
| Strain Kuutsalo-23 |  |  |  | 98.3% | 98.5% |
| Strain Haapasaari-18 |  |  |  |  | 98.6% |
| **Segment 3. NS3-like protein** |  |  |  |  |  |
| Strain Miass519 | 96.7% | 91.2% | 91.3% | 90.9% | 91.2% |
| Strain Miass527 |  | 90.7% | 91.1% | 90.5% | 91.0% |
| Strain H3 |  |  | 90.9% | 91.3% | 91.5% |
| Strain Kuutsalo-23 |  |  |  | 95.0% | 97.4% |
| Strain Haapasaari-18 |  |  |  |  | 94.5% |
| **Segment 4. VP2 and VP3** |  |  |  |  |  |
| Strain Miass519 | 98.6% | 90.5% | 90.8% | 90.7% | 91.2% |
| Strain Miass527 |  | 90.6% | 90.7% | 90.8% | 91.3% |
| Strain H3 |  |  | 90.6% | 90.3% | 91.1% |
| Strain Kuutsalo-23 |  |  |  | 94.7% | 98.0% |
| Strain Haapasaari-18 |  |  |  |  | 95.3% |

Overall, the full genomes of the Miass strains demonstrated >88% nucleotide identity to the other strains of ALSV known to date, with glycoprotein-coding segment 2 having the highest similarity and NS5-like protein coding segment 1 being the most divergent.

The two Miass strains were similar in their genome organization to each other and to known ALSV. Segments 1 and 3 were monocistronic. Segment 1 contained a flavivirus-NS5-like protein with polymerase and putative methyltransferase domains and a predicted signal peptide on the N-terminus of the protein. Segment 3 contained a flavivirus NS3-like protein with proposed functions of serine protease and RNA helicase (Figure 3). Both proteins had transmembrane regions corresponding to the previously-mapped strain H3. Segment 4 contained two non-flavivirus-related ORFs. There was a slippery sequence (G GTT TTT) and some evidence of an RNA-structured region (Supplementary file 7) and we believe that the VP3 ORF may be expressed via −1 ribosome frameshifting (Figure 3), as was previously shown for other ALSV strains. Regarding the VP1a and VP1b ORFs in segment 2, transmembrane regions and N-glycosylation sites were also in positions in the predicted amino acid sequences similar to the other ALSV strains (Wang et al., 2019a). Interestingly, we were not able to identify signal peptides in either VP2 or VP1a using TMHMM v.2.0 software.

**Figure 3.**
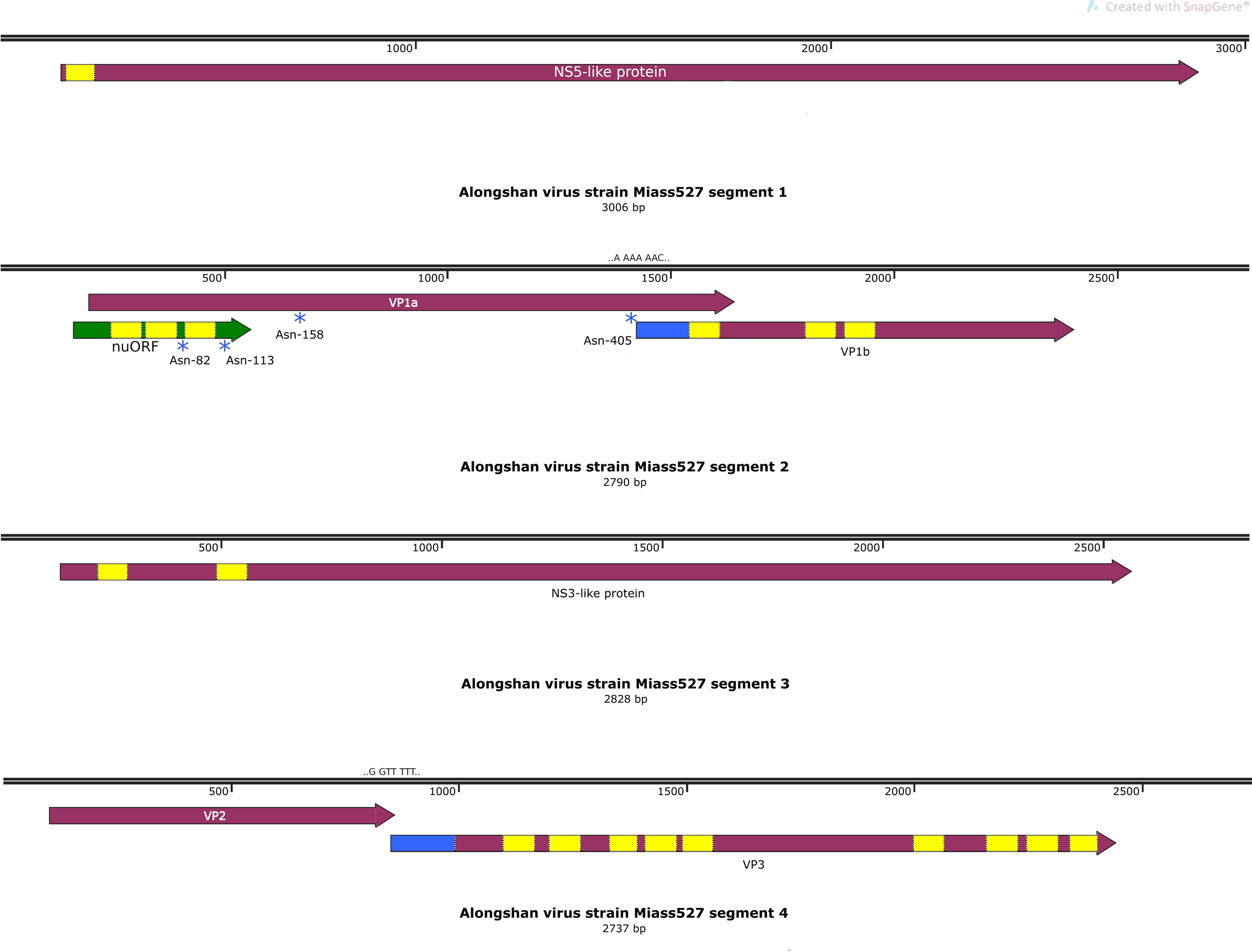
Genome of the Alongshan virus strain Miass527. Transmembrane regions are shown in yellow, the part of the VP1b protein derived from the proposed ribosomal frameshift is shown in blue, and the proposed novel protein in segment 2 is in green. Blue stars mark the putative N-glycosylation sites. Proposed ribosomal frameshift sites are in blue.

We performed a search of the functional elements in the VP1a region with the Synplot2 program (Firth, 2014). Using codon alignment of four previously-described ALSV strains (MH158416, MN107154, MN107158, MN095520) and the three strains sequenced in this study, we discovered two regions of high conservation in the VP1a ORF (Figure 4A). The first one corresponded to the 3`-terminal region of the ORF. We believe that such high conservation in this region was caused by an overlap with the VP1b ORF. We managed to find a slippery motif (A AAA AAC) followed by a spacer and an RNA structure (Supplementary file 8) in this region of the VP1a ORF that theoretically would allow VP1b to be translated via −1 ribosome frameshift (Figure 3).

**Figure 4.**
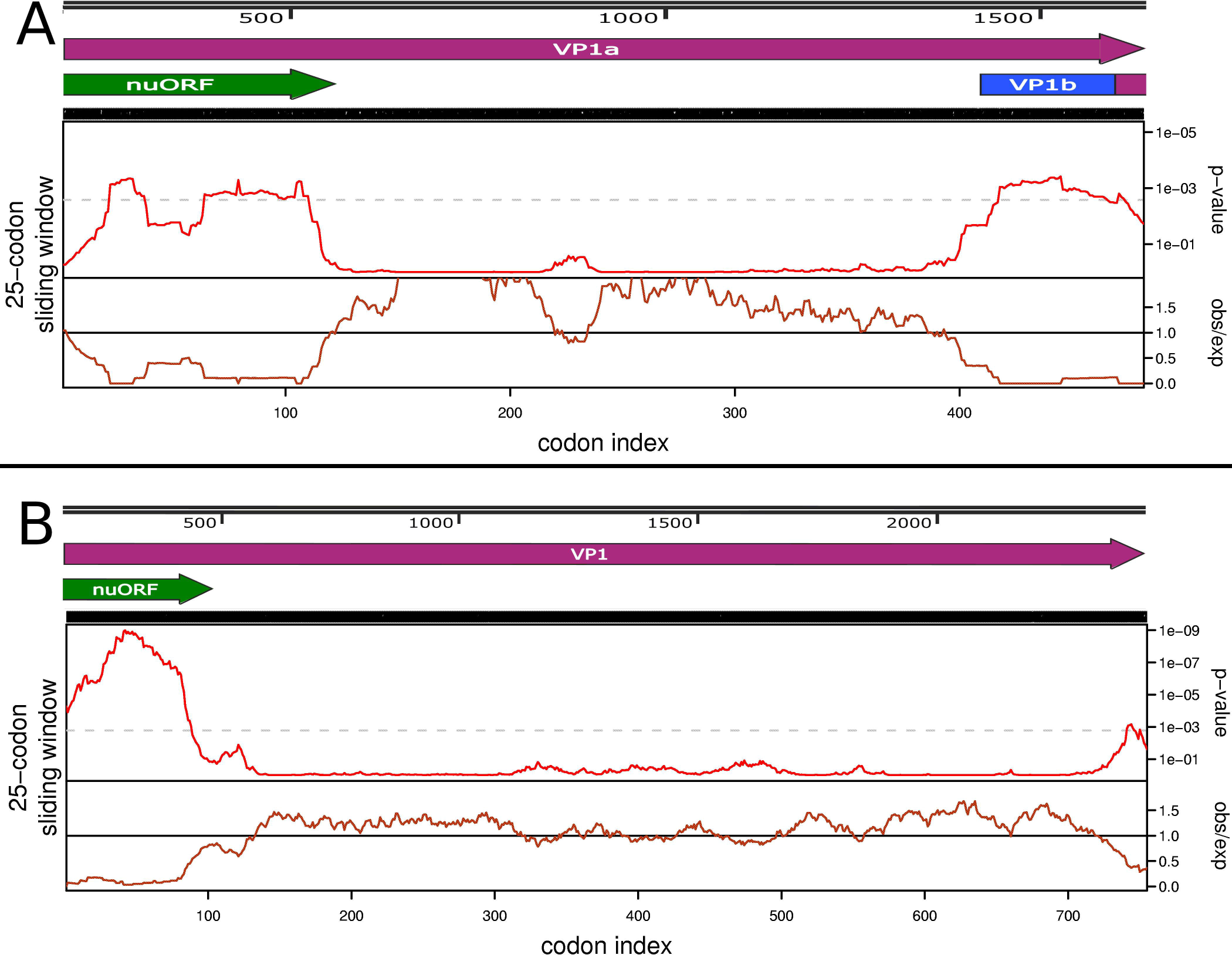
Assosiation of the nuORF with the region of high conservation in the Alongshan VP1a and Jingmen tick virus VP1 open reading frames. (A) Codon alignment of 7 full sequences of the Alongshan virus VP1a ORF was used in the analysis (Supplementary file 3). (B) Codon alignment of all 21 full sequences of the Jingmen tick virus VP1 ORF was used in the analysis (Supplementary file 3). In each section the top panel represents the virus genome map of the tested region. The middle panel depicts probability that the degree of ORF conservation within a 25-codon sliding window could be obtained under neutral evolution. The grey dashed line indicates p=0.005 significance (after correcting for multiple tests, where the number of tests is the length of a coding sequence divided by the window size). The bottom panel displays the relative amount of synonymous-site conservation at a 25-codon sliding window by showing the ratio of the observed number of synonymous substitutions to the expected number. The analysis was done using the Synplot2 program (Firth, 2014).

Additionally, Synplot2 allowed us to identify an unusually high conservation rate in the first ~110 codons of VP1a ORF in segment 2. We believe that this conservation was caused by an additional ORF that was found in this region. This novel upstream ORF (nuORF) is 399 nt long, is located upstream of the VP1a ORF and has a 365 nt overlap with it (Figure 4A).

Application of the tblastn algorithm using the protein sequence encoded by the Miass527 nuORF as an entry revealed homologs within the JMTV group (Supplementary file 9). Upon further inspection, all of the resultant sequences were shown to have a nuORF. Thus we decided to test known JMTV sequences for existence of the additional ORF upstream of the VP1 region with Synplot2.

We searched for a full-length segment 2 of JMTV using blast algorithm and strain SY84 JMTV full segment 2 sequence as an entry. Since relationships within the JMTV group are not yet decided and some representatives of this group have different names, we could not use a GenBank search. Sequences MN095532 and KX377514 were found to have unknown nucleotides in the middle of the segment and were not used in the analysis. For these sequences, nuORF was confirmed to be intact. We then constructed a phylogenetic tree (Supplementary file 10) that allowed us to identify three groups (Jingmen, Yanggou and Alongshan viruses) that should be analysed separately due to phylogenetic distance. All JMTV sequences with full segment 2 (21 entries) known to date (November 8, 2019) have a nuORF. We performed analysis of the JMTV segment 2 sequences (Figure 4B) and Synplot2 showed high conservation in the first ~100-codon region. This demonstrates that the region associated with nuORF is conserved more highly than would be expected under neutral evolution in both ALSV and JMTV sequences.

We also tested the Yanngou virus sequences and, although all of the full genome sequences had a nuORF, Synplot2 program did not find a region of higher-than-expected conservation in the first ~100 codons of Yanngou virus VP1a (Supplementary file 11). However, there are only three known full-length segment 2 sequences of Yanggou virus and the number of sequences is important to acquire statistics needed for a prediction.

Overall, the ALSV nuORF encoded 132 amino acids. According to the prediction, this protein may have an N-terminal signal sequence and three transmembrane regions with a C-terminal tail, thus making the nuORF product a small membrane protein (Figure 5A).

**Figure 5.**
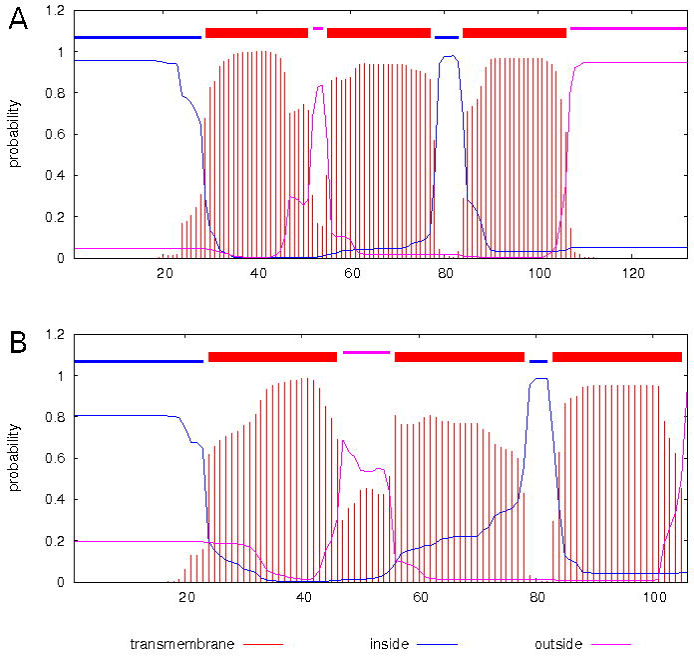
Transmembrane domains of the proposed nuORF product. of (A) Alongshan virus strain Miass527, (B) Jingmen tick virus strain SY84. The analysis was done using TMHMM Server v. 2.0 program (Krogh et al., 2001).

The predicted protein encoded by the nuORF of JMTVs was shorter (~106 aa) and had ~50% identity to the ALSV nuORF product. It had an overall structure similar to that of ALSV, with a proposed N-terminal signal sequence and three predicted transmembrane regions, but, in contrast to the ALSV nuORF, it had no C-terminal tail (Figure 5B).

## Discussion

Due to the development of molecular virology, high-throughput sequencing and viral metagenomics, many new flavi-like viruses have been discovered in recent decades (Qin et al., 2014; Shi et al., 2016; Villa et al., 2017). A large number of flavi-like viruses have been detected in or isolated from arthropods (Huhtamo et al., 2014, 2009; Shi et al., 2016), but their ability to cause infection and disease in vertebrates and their medical significance are not known.

The JMTV group is a recently-discovered group of segmented ssRNA(+) viruses that infect different species of invertebrates, including insects, ticks and nematodes (Callister et al., 2008; Jia et al., 2019; Kuivanen et al., 2019; Qin et al., 2014; Shi et al., 2016; Tetteh et al., 1999; Webster et al., 2015). Moreover, some representatives of the new flavi-like viruses were detected in patients who had other severe infectious diseases (Emmerich et al., 2018), and these viruses might themselves cause illness in humans (Jia et al., 2019; Wang et al., 2019a). There is still little information about the replication and overall coding strategy of this novel virus group.

ALSV is a representative of the JMTV group. It was first isolated in China from a patient suffering fever of unknown aetiology (Wang et al., 2019a). Subsequently, the virus was detected in *I. persulcatus* ticks and *Culex tritaeniorhynchus* and *Anopheles yatsushiroensis* mosquitoes in China (Wang et al., 2019a) and in *I. ricinus* ticks in Finland (Kuivanen et al., 2019) and France (Temmam et al., 2019). It was shown that ALSV is able to infect mammals and cause illness in humans (with symptoms including headache, fever, fatigue and depression) (Wang et al., 2019a, 2019b). Thus, additional isolates of this virus from new geographical locations will be important for understanding its distribution, as well as for the local healthcare systems. New genome data could also provide additional information on the structural organization of the ALSV genome.

We isolated three strains of ALSV from *I. persulcatus* ticks collected in Chelyabinsk region in the south of European Russia, and in the Republic of Karelia in north-west Russia. We sequenced two full genomes of the new ALSV strains Miass527 and Miass519 from Chelyabinsk region and obtained partial sequences of segment 2 for the ALSV strain Galozero-14-T20426 from the Republic of Karelia. Although these three strains were isolated from territories distant from each other, phylogenetic analysis of the segment 2 sequences showed that they form a single monophyletic group with the strain H3 isolated in China.

The segments 1, 3 and 4 of the strains Miass519 and Miass527 were more similar to each other than to previously described ALSV strains. Segment 2 of the strain Miass527 was more similar to the strain H3 than to Miass519, while segment 2 of the strain Miass519 was closer to the isolate JMTV/I.ricinus/France then to Miass527. This data suggests that reassortment of genome segments within ALSV could occur. Similar observations were made earlier for JMTV (Qin et al., 2014).

Chinese strains of ALSV were isolated from mammalian blood samples using Vero (African green monkey) cells (Wang et al., 2019a, 2019b) while the isolation of the virus detected in Finnish ticks failed in Vero and two human cell lines (Kuivanen et al., 2019). The Chinese JMTV strain isolated from *A. javanense* ticks replicated successfully in the *R. microplus* embryo-derived tick cell line BME/CTVM23 but failed to grow in Vero, DH82 (canine monocyte-macrophage-derived cells) and C6/36 (*Aedes albopictus* mosquito) cells (Jia et al., 2019). We isolated three tick-derived strains of ALSV using the tick cell line IRE/CTVM19; our results, together with those of the previous studies reporting successful *in vitro* isolation and continuous propagation of JMTV group viruses (Jia et al., 2019; Wang et al., 2019a, 2019b), suggest the possibility that mammal-derived viruses will more readily infect mammalian cells, while tick-derived viruses may be more infective for tick cells. However, additional isolations of JMTV group viruses will be required to test this theory and the possible underlying genetic causes. For the first time we showed that the ALSV strain Miass527 can persist in tick cells for at least three years. It was previously shown that the tick cell line IRE/CTVM19 may harbour an unknown rhabdovirus (Bell-Sakyi and Attoui, 2016). Our experiment confirmed the presence of a rhabdovirus and suggests that it does not interfere with the persistence and active reproduction of ALSV in IRE/CTVM19 cells. Persistent infection in tick cells and ticks could affect virus properties, as was previously shown for tick-borne encephalitis virus (Belova et al., 2017).

Previously, it was shown that the virion sizes of JMTV group representatives ranged from 30-35 nm to 70-100 nm (Ladner et al., 2016; Qin et al., 2014; Wang et al., 2019a). ALSV virions were reported to be enveloped spherical (or nearly spherical) particles with a diameter of approximately 80-100 nm (Wang et al., 2019a). The size of virions of our strain Miass527 was 40.5±3.7 nm. In transmission electron microscopy, some of the viral particles had electron-dense cores, and could hypothetically be defective virions without genomic RNA (Korboukh et al., 2014). In addition, we found smaller particles with a diameter of 13.1±2.1 nm, which had either electron-translucent or electron-dense cores. Presumably, they could represent either: a) glycoprotein aggregates, similar to the “classic” flavivirus nonstructural protein 1, which is secreted from infected cells (Alcalá et al., 2016; Muller et al., 2012), b) protein structures appearing after long-term virus persistence, c) virions with incomplete genome, or d) virions of an unknown virus or other unknown structures present in IRE/CTVM19 cells. In the equivalent gradient fraction of uninfected cells, we found only particles with a diameter of 15.7±1.8 nm, i.e., slightly larger than the small particles that we saw in the same fraction of Miass527-infected cells. In any case, the particles were smaller than the smallest parvovirus virions with a diameter of 18-26 nm (Mi et al., 2019). Further investigations are needed to clarify the nature of these particles.

Although the strain Miass527 of ALSV did not differ in overall genome structure from the previously isolated ALSV (Wang et al., 2019a), the newly-obtained data allowed us to propose a novel protein encoded by both ALSV and JMTV and, probably, by some other members of the JMTV group. The nuORF, proposed here, is conserved in all the full segment 2 sequences of ALSV, JMTV and Yanngou virus available at the time of writing. Considering that ALSV and JMTV are somewhat distantly related (around 70% nucleotide identity and around 80% amino acid identity in the NS5-like protein), it is likely that the nuORF indeed encodes a functional virus protein. Nevertheless, further studies are needed to confirm its expression and function during the virus life cycle.

Interestingly, an ORF of a more distant member of the JMTV group, the mosquito-borne Guaico Culex virus (Ladner et al., 2016), was discovered in the same genome region. It also encodes a small protein, but with no amino acid identity to the nuORF. A mass spectrometry analysis of the Guaico Culex virus virion proteins did not identify any peptides that belong to this putative ORF, thus making its product (if it is indeed expressed) a nonstructural protein (Ladner et al., 2016).

If the nuORF product protein is indeed functional at some point during ALSV replication, it would most likely be a minor membrane protein. Such proteins are present in a variety of RNA viruses and perform different functions. For example, in flaviviruses, small membrane proteins NS2B and NS4A (with sizes similar to the nuORF) perform multiple functions, ranging from enzyme cofactors (Lindenbach and Rice, 2003; Shiryaev et al., 2009) to participation in particle formation (Li et al., 2016; Muñoz-Jordan et al., 2003) and protection against the cellular antiviral response. Influenza A M2 protein acts as a proton-selective ion channel (DiMaio, 2014), whereas *Coronavirus* E protein acts as a viroporin and plays a role in induction of the regulation of the unfolded protein response (Schoeman and Fielding, 2019). Nevertheless, confirmation of the nuORF expression and understanding of its possible functions should be subjects for further studies.

In conclusion, we isolated three strains of ALSV from *I. persulcatus* ticks in Russia. Although these strains were isolated from territories far from each other, in phylogenetic analysis they formed a single monophyletic group with the virus isolated in China. Our study confirmed the value of tick cell lines in virus isolation and maintenance of persistent infection. Availability of these novel ALSV strains has enabled better understanding of the phylogenetic relationships within the JMTV group, and will facilitate further study of the origins, structure, replication, epidemiology, transmission, vertebrate hosts and medical and veterinary significance of these enigmatic segmented RNA viruses.

## Materials and Methods

### Collection and Processing of Ticks

In 2014, from the beginning of May to the middle of June, unfed adult ticks were collected by flagging from vegetation in the Republic of Karelia and Chelyabinsk region (Supplementary file 1). Ticks were identified using taxonomic keys (Filippova, 1997, 1977). Adult ticks were homogenised in pools of 3-5 individuals according to species composition, location, and route of collection using the laboratory homogeniser TissueLyser II (QIAGEN, Germany) in Medium 199 supplemented with Earle’s salts (FSBSI Chumakov FSC R&D IBP RAS, Russia) and antibiotics. All homogenates were tested for the presence of flavivirus RNA using a heminested RT-PCR and primers targeting a conserved region of the flavivirus NS5 gene, as described earlier (Scaramozzino et al., 2001).

### Infection of Tick Cell Line

We used a cell line derived from embryos of the tick *I. ricinus* − IRE/CTVM19 (Bell-Sakyi et al., 2007) provided by the Tick Cell Biobank. The tick cell line was maintained at 28 °С in L-15 (Leibovitz) medium (FSBSI Chumakov FSC R&D IBP RAS, Russia) supplemented with 10% tryptose phosphate broth, 20% fetal bovine serum, 2 mM L-glutamine and antibiotics as described earlier (Weisheit et al., 2015). Prior to infection, IRE/CTVM19 cells were seeded in flat-sided culture tubes (Nunc, ThermoFisher Scientific) in 2.2 ml of complete medium and incubated at 28 °C. A week later, cells were infected by adding 200 μl of virus-containing material (unfiltered tick homogenate or culture supernate) and incubated at 28 °С. Medium was changed at fortnightly intervals by removal and replacement of 1.5 ml; the spent medium was used to harvest the virus as described below. Occasional subcultures were carried out at intervals between 1 month and >1 year by adding 2.2ml of fresh medium, resuspending the tick cells and transferring half of the cell suspension to a new culture tube.

For transmission electron microscopy and high-throughput sequencing, infected cell culture supernate (spent medium) was clarified by centrifugation at 10,000 rpm for 30 min at 4 °C using an SW-28 rotor in an Optima L-90K Ultracentrifuge (Beckman Coulter, USA) and was then ultracentrifuged at 25,000 rpm for 6 h at 4 °C using the same rotor. Uninfected cell culture supernate was prepared similarly for transmission electron microscopy.

### Transmission Electron Microscopy

The pellet obtained following ultracentrifugation of culture supernate was dissolved overnight in borate saline buffer (0.05M H_3_BO_4_, 0.12M NaCl, 0.24M NaOH), pH 9.0 (BSB), at 4 °C. A portion of the dissolved pellet was applied to a sucrose density gradient (15-60%) and centrifuged at 30,000 rpm for 4 h at 4°C using an SW-40 rotor in the Optima L-90K Ultracentrifuge (Beckman Coulter, USA). To identify resultant density gradient fractions containing virus particles, aliquots of each fraction were subjected to RT-PCR and sequencing of amplified products as described below.

The specimens after ultracentrifugation were adsorbed onto Formvar film attached to 200-mesh nickel EM grids and contrasted with 2% uranyl acetate (Nikitin et al., 2015). The grids were exposed to UV light for one hour to inactivate the virus, air-dried and examined in a JEOL JEM-1400 transmission electron microscope (JEOL, Tokyo, Japan) operated at 80 kV. Images were acquired with an Olympus Quemesa digital camera using iTEM software (Olympus Soft Imaging Solutions GmbH, Munster, Germany).

### Reverse-transcriptase PCR (RT-PCR) and Sequencing of Amplified Products

Viral RNA from tick suspensions and culture supernate of infected cells was isolated with TRI Reagent LS (Sigma-Aldrich, USA) according to the manufacturer’s protocol. Reverse transcription was performed with random hexamer primers (R6) and M-MLV reverse transcriptase (Promega, Madison, WI) according to the manufacturer’s protocol. Viral genomic cDNA was amplified by PCR using specific primers MiassF and MiassR (Supplementary file 2). Sequencing was carried out in both directions directly from PCR-amplified DNA on the ABI PRISM 3730 (Applied Biosystems) sequencer using ABI PRISM® BigDye™ Terminator v. 3.1. Genomic sequences were assembled using SeqMan software (DNAstar, USA).

### High-Throughput Sequencing

Total RNA from an ultracentrifugation pellet was fragmented and reverse-transcribed into cDNA with R6 primers using RevertAid reverse transcriptase (ThermoFisher Scientific) followed by second strand synthesis with the NEBNext Ultra II Non-Directional RNA Second Strand Synthesis Module (New England Biolabs Inc. USA). Resultant double-stranded DNA was purified using Ampure XP (Beckman Coulter) and subjected as an input for library preparation process using the NEBNext® Fast DNA Library Prep Set for Ion Torrent™ (New England Biolabs Inc. USA) following the manufacturer’s instructions. The resultant DNA library was quantified with the Ion Library TaqMan™ Quantitation Kit (ThermoFisher Scientific) followed by templating on the Ion Chef instrument (ThermoFisher Scientific) and sequencing on the Ion S5XL instrument with the viral library constituting a part of the Ion 530 Chip. Raw reads were quality-controlled using FaQCs v2.09 (Lo and Chain, 2014) and assembled into contigs using SPAdes v3.13.0 (Bankevich et al., 2012) with iontorrent flag. The resultant contigs were screened for viral sequences using Blastn algorithm in BLAST v2.9.0+ with nt database and contigs corresponding to virus were extracted for further investigation.

### Phylogenetic Analysis

RNA sequences of all published strains of ALSV, representatives of the JMTV group and the three strains described in this article were used in the phylogenetic analysis. The nucleotide sequences of the genome coding region of segment 2 were aligned using Clustal-X 2.0.11. Phylogenetic analysis was conducted using the neighbour-joining method using MEGA 6.0 with 1000 bootstrap replications.

### Bioinformatics tools used for discovery of overlapping elements

For the analysis we used sequences of the full segment 2 of ALSV, JMTV and Yanggou virus extracted from GenBank (08.11.19), as well as sequences of the ALSV strains obtained in the present study (Supplementary file 9). Sequences were divided into three groups (named “JMTV”, “ALSV” and Yanggou virus) based on their positions in the phylogenetic tree (Supplementary file 10). Sequences obtained in the current study were included in the ALSV alignment.

Analyses of Yanggou virus, ALSV and JMTV were performed separately, because of a low level of similarity between them. Nucleotide sequences of the ORF VP1a (in the case of ALSV and Yanggou virus) and VP1 (in the case of JMTV) were extracted from the full segments. The resultant sequences were codon-aligned using the online tool «Codon Alignment v2.1.0» (available at https://hcv.lanl.gov/content/sequence/CodonAlign/codonalign.html) with default parameters. The resultant codon alignments were used as an input for the Synplot2 (Firth, 2014) online tool (available at http://guinevere.otago.ac.nz/cgi-bin/aef/synplot_c.pl) v. 2014-12-05/03:10:08 with default parameters (no reference sequence, n=12 (sliding window of 25 codons)).

### Genome annotation and visualization

Putative N-glycosylation sites were predicted using the NetNGlyc 1.0 Server (available online at http://www.cbs.dtu.dk/services/NetNGlyc/) with a standard set of parameters (Gupta et al., 2004). Transmembrane region prediction was done using the TMHMM Server v. 2.0 (available online at http://www.cbs.dtu.dk/services/TMHMM/) (Krogh et al., 2001). Aligned ALSV sequences were examined with the pAliKiss web server (build in June 25 2015, available https://bibiserv.cebitec.uni-bielefeld.de/palikiss) with default parameters (Janssen and Giegerich, 2015) to find possible conserved RNA structures downstream of putative frameshift sites. ORFs, transmembrane regions and N-glycosylation sites were mapped on the genome using SnapGene Viewer v. 4.3.10. Further image editing was performed in the GNU Image manipulation program v. 2.10 software.

## Acknowledgments

Tick collection by Sergey V. Bugmyrin and Liubov A. Bespyatova is supported by the state order (project № 0218-2019-0075). Lesley Bell-Sakyi is supported by the United Kingdom Biotechnology and Biological Sciences Research Council grant number BB/P024270/1.

