## Supplementary files for "First isolation and characterisation of Alongshan virus in Russia"

**Supplementary file 1. Regions of Russia where tick collection was carried out.** In 2014 from the beginning of May to the middle of June, adult *Ixodes persulcatus* ticks were collected by flagging from vegetation in the Republic of Karelia and the Chelyabinsk region in Russian Federation.

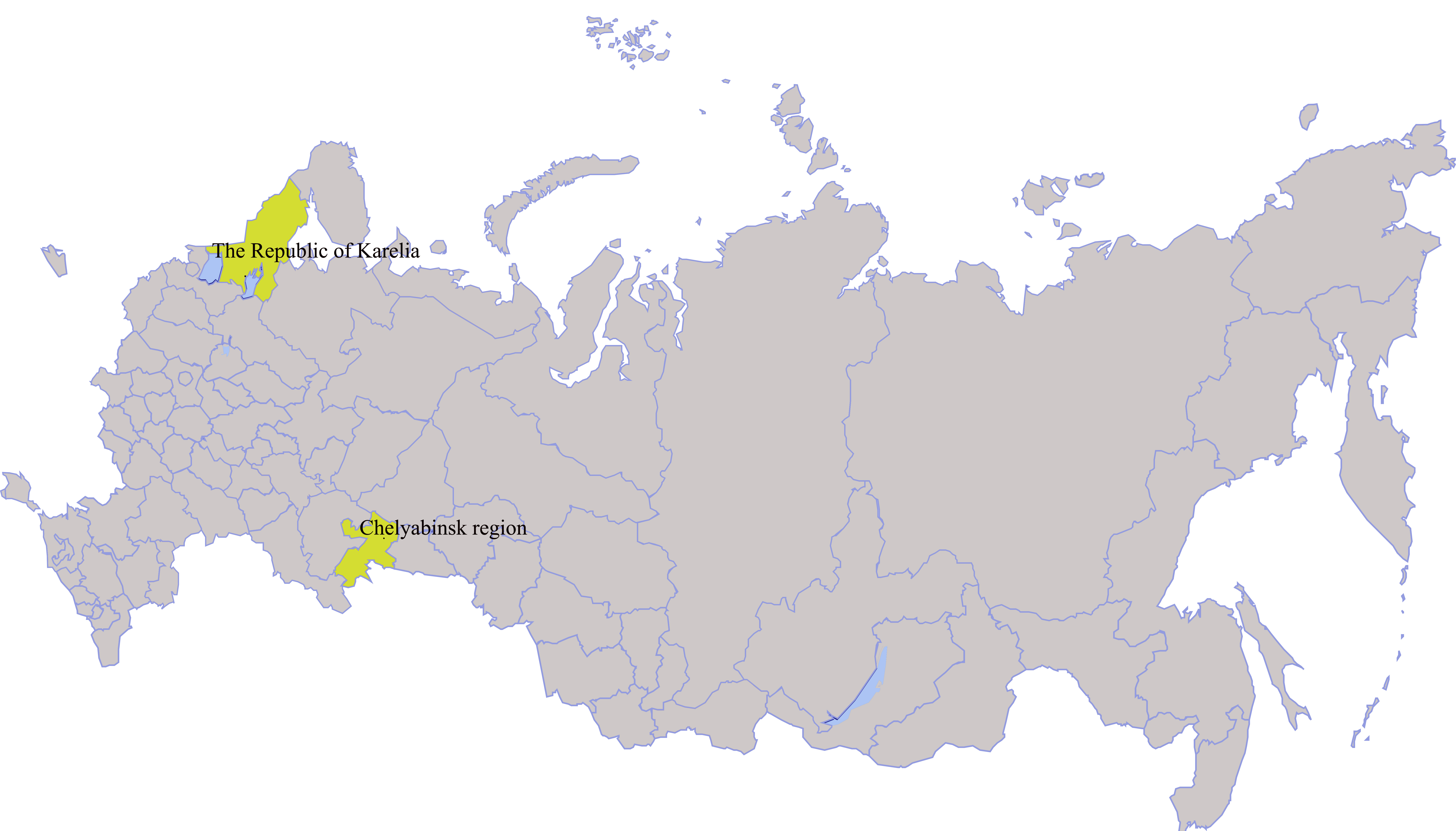

**Supplementary file 2. Specific primers for amplification of genome segments 1 and 2 of Alongshan virus and IRE/CTVM19-associated rhabdovirus.**

| Name of primers | Nucleotide sequence | Primer direction | Genome locus | Amplicon size, bp | Temperature, ^0^C |
| --- | --- | --- | --- | --- | --- |
| MiassF | GGTACACGGACCTGGGATCCTATTG | Forward | segment 1 | 825 |  |
| MiassR | TCTCTGACTCCTGTTCTAATC | Reverse | segment 1 |  |  |
| JMun1S | TTAAAARCGGCCAGCCTTNRYTGCAAGTGCA | Forward | segment 2 | 1800 |  |
| Miass_gly_1R | ACCAGGTTGGTCAAGGCAAT | Reverse | segment 2 |  |  |
| Miass_gly_3F | TGGATCAGCTCACACCACAC | Forward | segment 2 | 333 |  |
| Miass_gly_3R | TCACCGTCACAGTGGAATGG | Reverse | segment 2 |  |  |
| Rhabdo _L_1F | GGGTTTGTGGTTAATTTGTC | Forward |  | 392 |  |
| Rhabdo_L_1R | AGTGAGGACTGGATAAAAGA | Reverse |  |  |  |

**Supplementary file 3. Electrophoresis of PCR fragments of sucrose density gradient fractions of Alongshan virus-infected IRE/CTVM19 cells.**

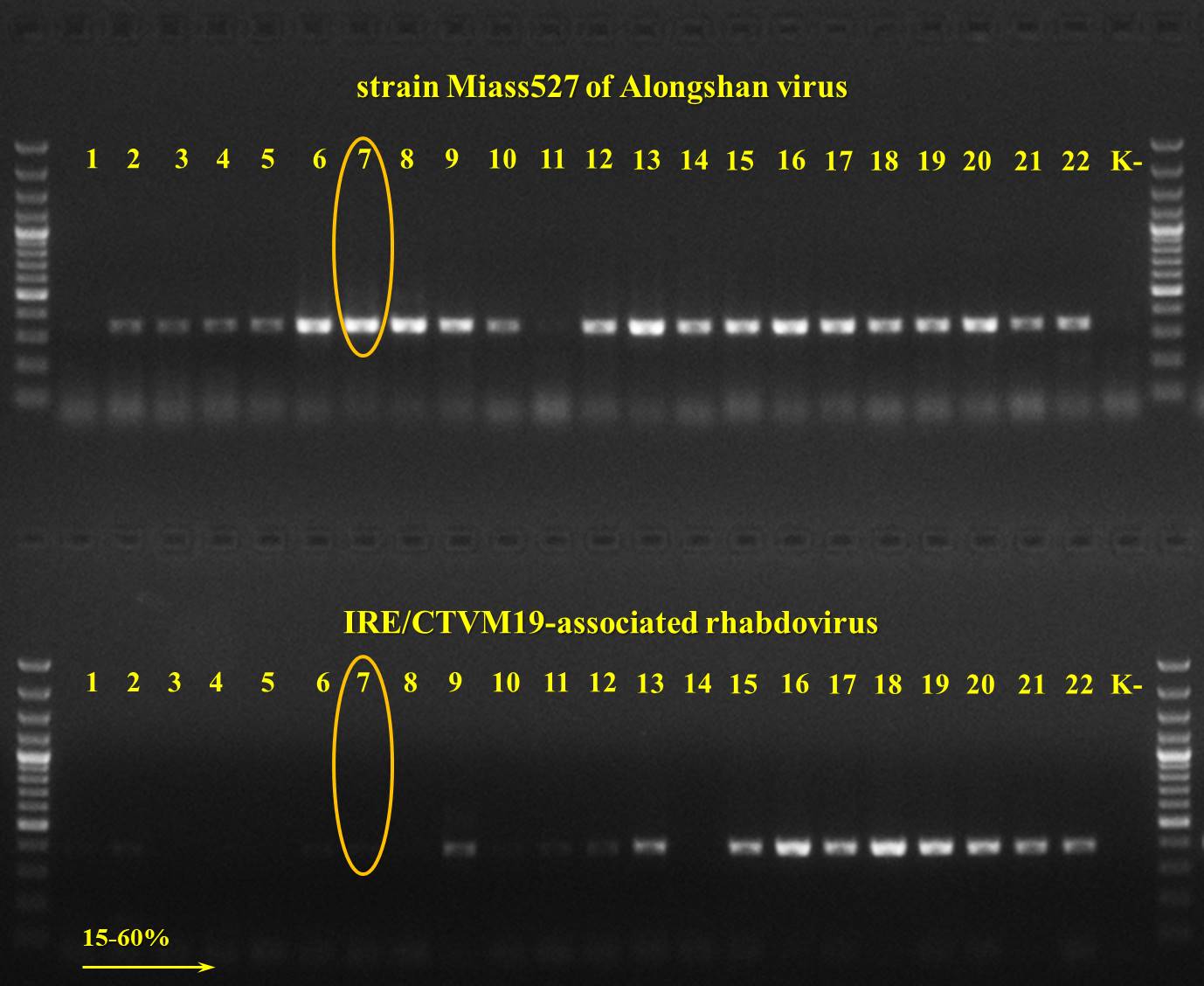

We separated Alongshan virus (ALSV) strain Miass527 and IRE/CTVM19-associated rhabdovirus by sucrose density gradient centrifugation to identify the fraction with the largest amount of ALSV RNA. Viral genomic cDNA was amplified by PCR using specific primers for ALSV (Miass_gly_3F and Miass_gly_3R) and IRE/CTVM19-associated rhabdovirus (Rhabdo _L_1F and Rhabdo_L_1R). Fraction #7 was selected for transmission electron microscopy.

**Supplementary file 4. The range of virion size** **detected in transmission electron microscopy of strain Miass527 of Alongshan virus.**

Most of the virions were spherical particles with a diameter of 40.5 ± 3.7 nm.

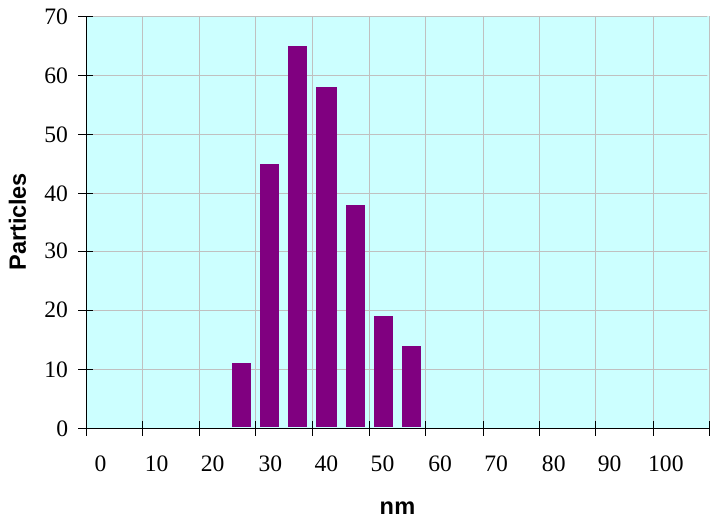

**Supplementary file 5. The size of the small spherical particles detected in transmission electron microscopy of strain Miass527 of Alongshan virus.**

The small spherical particles had a diameter of 13.1 ± 2.1 nm.

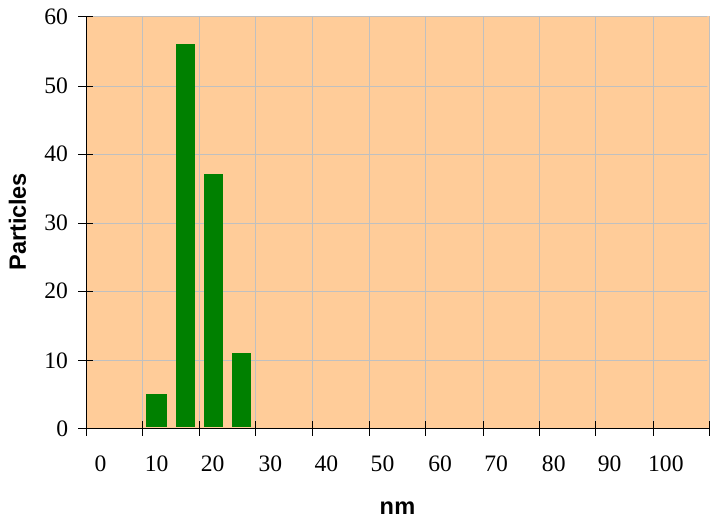

**Supplementary file 6. The size of the small spherical particles detected in transmission electron microscopy of ultracentrifuged supernate from uninfected IRE/CTVM19 cells.**

The small spherical particles had a diameter of 15.7 ± 1.76 nm.

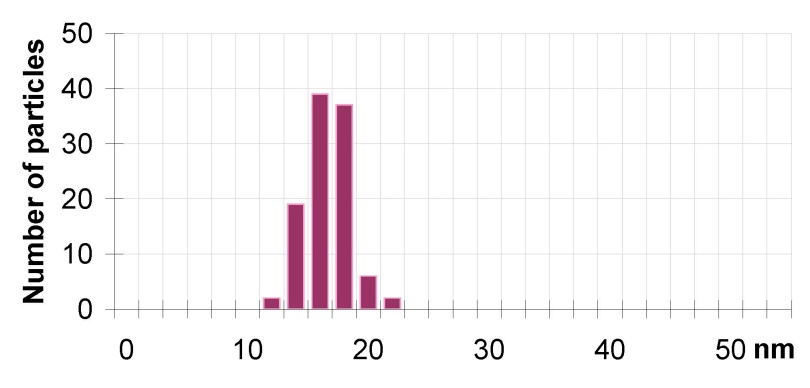

**Supplementary file 7. Predicted RNA structure downstream of the frameshift region in segment 4 of Alongshan viruses (ALSV).**

The frameshift site is colored teal (turquoise). Red, green and grey coloring represents paired groups of nucleotides in the structure that may cause frameshifts. Structure was predicted with the pAliKiss algorithm (Stefan Janssen and Robert Giegerich The RNA shapes studio, Bioinformatics, 2015) using alignment of the 201 nt region of ALSV strains H3, Miass527, Miass519, Kuutsalo-23 and Haapasaari-18 as an entry.

(-75.94 = -74.90 + -1.04)

1 gguuuuucaguaggggaggggaauacccagccaccccccucccgaaccgacaggaagccugucggagaagggaagagcuggauaccgaacugggcugaagugagcagggacuacuugguuagacucggaaagaaggccgcacucaugggagagggcuacgacaucauccuugaugaggcuagggauuuguucccaaccucc 201

.........[[..{{{{{{{{.]]..<<<<<....}}}}}}}}...((((((((...))))))))...........>>>>>...[[[[....{{{{....(((.((((.....)))).)))...]]]]..<<<.((((...(((....)))..))))....(((((....))))).}}}}.((((.....))))....>>>

**Supplementary file 8. Predicted RNA structure downstream of the frameshift region in segment 2 of Alongshan viruses (ALSV).**

The frameshift site is colored teal (turquoise). Red, green, grey, purple and blue coloring represent paired groups of nucleotides in the structure that may cause frameshifts. Structure was predicted with the pAliKiss algorithm (Janssen, Stefan and Giegerich, Robert The RNA shapes studio, Bioinformatics, 2015) using alignment of the 201 nt region (downstream to the proposed frameshift site) of ALSV strains H3, Miass527, Miass519, Kuutsalo-23 and Haapasaari-18 as an entry.

1 aaaaaacgucaucagaccguccaugaggcuuuuaccgauuacacccccgugccaaagcauugguacgacuggcuuucgagcugguuuggccacaucacgggagguaucgagagggcuuaucggaucaucucaugggcggucgaguuuguuacuagcaacgcuccaacugugcuuguggucaucaucauggccacaccccuc 201

(-75.50 = -74.66 + -0.84) .......[[[....{{{{{{{{{{{{]]]......((((...(((((((((.........((((.(((((((((....)))))).)))))))...)))))).))))))).<<<<<..............}}}}}}}}}}}}(((((((((...))))).))))..........(((((((((....))))))))).>>>>>

**Supplementary file 9. List of Jingmen tick virus group sequences used for the discovery of the conservative elements within segment 2.**

| Accession number | Virus name (according to GenBank) | Used for analysis |
| --- | --- | --- |
| MH158416 | Alongshan virus strain H3 | ALSV ^1^ |
| MN107154 | Alongshan virus strain Kuutsalo-23 | ALSV |
| MN107158 | Alongshan virus strain Haapasaari-18 | ALSV |
| MN095520 | Jingmen tick virus isolate JMTV/I.ricinus/France | ALSV |
| MH688530 | Yanggou tick virus strain YG | Yanggou virus ^2^ |
| MH688533 | Yanggou tick virus strain 16-T2 | Yanggou virus |
| MH688537 | Yanggou tick virus strain 17-L1 | Yanggou virus |
| KJ001580 | Jingmen Tick Virus isolate SY84 | JMTV^3^ |
| KY523073 | Mogiana tick virus isolate MGTV/V4/11 | JMTV |
| MG703254 | Amblyomma virus GXTV108 | JMTV |
| MH133315 | Jingmen tick virus isolate Kosovo 2013-17-266 | JMTV |
| MH133319 | Jingmen tick virus isolate Kosovo 2014-C-K14-1C | JMTV |
| MH133323 | Jingmen tick virus isolate Kosovo 2015-A-K15-1A | JMTV |
| MH155890 | Jingmen tick virus isolate JTMV_1 | JMTV |
| MH155894 | Jingmen tick virus isolate JTMV_3 | JMTV |
| MH155905 | Jingmen tick virus isolate JTMV_100 | JMTV |
| MH814978 | Rhipicephalus associated flavi-like virus isolate YNTV4 | JMTV |
| MK174244 | Jingmen tick virus isolate XJ58 | JMTV |
| MK174245 | Jingmen tick virus isolate XJ61 | JMTV |
| MK174246 | Jingmen tick virus isolate XJ77 | JMTV |
| MK174247 | Jingmen tick virus isolate XJ155 | JMTV |
| MK174248 | Jingmen tick virus isolate XJ335 | JMTV |
| MK174249 | Jingmen tick virus isolate XJ363 | JMTV |
| MK673134 | Kindia tick virus isolate KITV/2017/1 | JMTV |
| MN025513 | Jingmen tick virus isolate TTP-Pool-3b | JMTV |
| MN025517 | Jingmen tick virus isolate TTP-Pool-19 | JMTV |
| MN095524 | Jingmen tick virus isolate JMTV/Rh.microplus/Am.variegatum/French Antilles | JMTV |
| MN095528 | Jingmen tick virus isolate JMTV/Am.testudinarium/Lao PDR | JMTV |
| MN095532 | Jingmen tick virus isolate JMTV/Pteropus lylei/Cambodia | None^4^ |
| KX377514 | Jingmen tick virus strain RC27 | None |

^1^ – VP1a ORF was used in the codon alignment of Alongshan viruses (ALSV)

^2^ – VP1a ORF was used in the codon alignment of Yanggou viruses

^3^ – VP1 ORF was used in the alignment of Jingmen tick virus (JMTV)

^4^ – sequence was not used in any of the alignments above due to large number of unidentified nucleotides. nuORF was confirmed to be intact.

**Supplementary file 10. Phylogenetic tree of all full segment 2 sequences of the Jingmen tick virus group.**

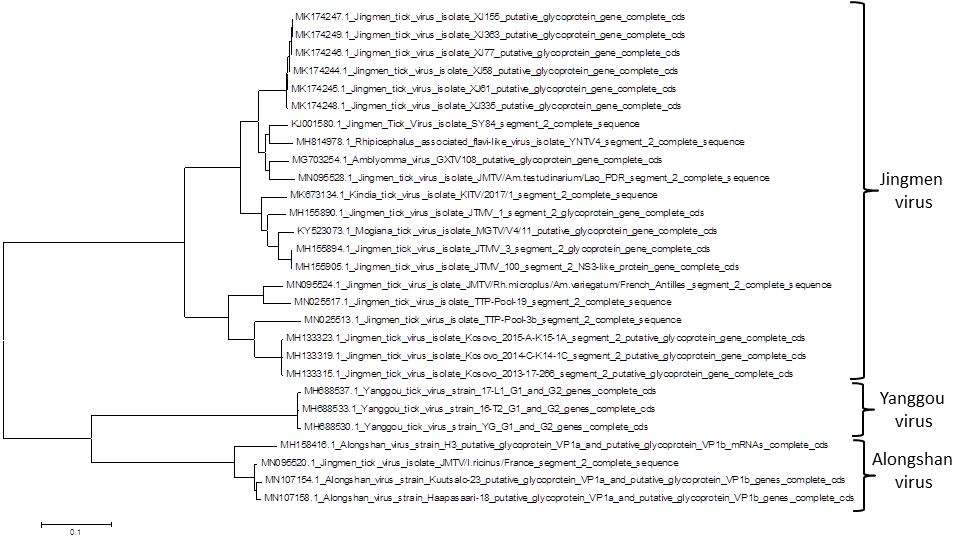

The evolutionary history was inferred using the Maximum Likelihood method based on the Tamura-Nei model (Tamura K. and Nei M. (1993). Estimation of the number of nucleotide substitutions in the control region of mitochondrial DNA in humans and chimpanzees. Molecular Biology and Evolution 10:512-526.). The tree with the highest log likelihood (-16975.0885) is shown. Initial tree(s) for the heuristic search were obtained automatically by applying Neighbor-Join and BioNJ algorithms to a matrix of pairwise distances estimated using the Maximum Composite Likelihood (MCL) approach, and then selecting the topology with superior log likelihood value. The tree is drawn to scale, with branch lengths measured in the number of substitutions per site. The analysis involved 28 nucleotide sequences. All positions containing gaps and missing data were eliminated. There were a total of 2254 positions in the final dataset. Evolutionary analyses were conducted in MEGA5 [(Tamura K., Peterson D., Peterson N., Stecher G., Nei M., and Kumar S. (2011). MEGA5: Molecular Evolutionary Genetics Analysis using Maximum Likelihood, Evolutionary Distance, and Maximum Parsimony Methods. Molecular Biology and Evolution 28: 2731-2739.).

**Supplementary file 11. Synonymous site conservation analysis of the Yanngou virus VP1a ORF**.

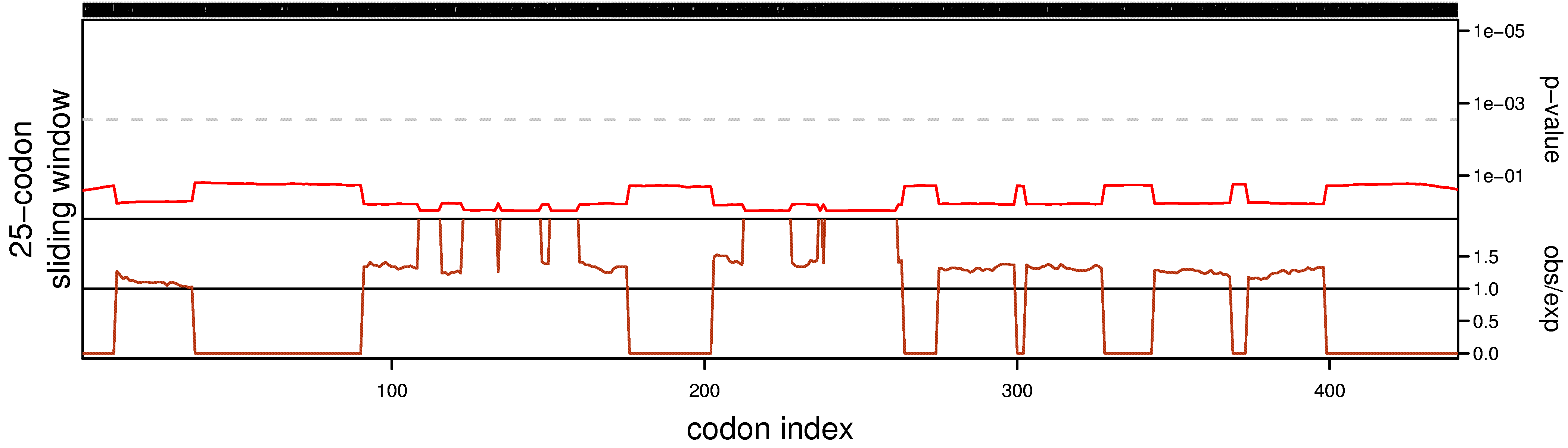

The top panel depicts the probability that the degree of ORF conservation within a 25-codon sliding window could be obtained under neutral evolution. The grey dashed line indicates p=0.005 significance (after correcting for multiple tests, where the number of tests is the length of a coding sequence divided by the window size). The bottom panel displays the relative amount of synonymous-site conservation at a 25-codon sliding window by showing the ratio of the observed number of synonymous substitutions to the expected number.

The analysis was done using Synplot2 program (Firth, 2014).
